# CSF pressure in fetal mice *in utero*: External factors pressurize the intraventricular space

**DOI:** 10.1101/2024.09.08.611845

**Authors:** Koichiro Tsujikawa, Reina Muramatsu, Takaki Miyata

**Author notes:** Corresponding author: Koichiro Tsujikawa, Takaki Miyata Department of Anatomy and Cell Biology, Nagoya University Graduate School of Medicine, 65 Tsurumai, Showa, Nagoya, 466-8550, Japan.

## Abstract

Previous experiments inducing leakage of embryonic cerebrospinal fluid (CSF) suggest the necessity of intraventricular CSF pressure (P_CSF_) for brain morphogenesis. Nevertheless, how embryonic P_CSF_ occurs is unclear, especially *in utero*. Using a Landis water manometer, we measured P_CSF_ in fetal mice isolated from the amniotic cavity (P_CSF-ISO_). At embryonic day (E) 13, P_CSF-ISO_ was 82.7 Pa. Intraventricular injections of ≥2 μl of saline elevated P_CSF-ISO_ by ∼30%. Intraventricularly injecting inhibitors of CSF secretion decreased P_CSF-ISO_ by ∼30%. Shh-mediated cerebral-wall expansion and the resulting ventricular narrowing did not significantly increase P_CSF-ISO_. Removal of the brain-surrounding contractile tissues decreased P_CSF-ISO_ by 80-90%. The intraamniotic pressure measured *in utero* (P_AF-IU_) was 1030.7 Pa. Our direct measurement revealed that the P_CSF_ *in utero* (P_CSF-IU_) was 1076.4 Pa, confirming the susceptibility of P_CSF_ to external factors. Subsequent P_CSF_ measurements under hydrostatic pressure loading suggested that P_CSF-IU_ = P_CSF-ISO_ + P_AF-IU_, a relationship further used to estimate P_CSF-IU_ at other ages when direct measurement was not possible. The estimated P_CSF-IU_ decreased almost constantly from E10 to E16 (2000 ➔ 500 Pa).

## Introduction

In developing embryos, the brain ventricles filled with embryonic cerebrospinal fluid (CSF) are surrounded by the primordial brain wall, together called the brain vesicles (Gato and Desmond, 2009; Chau et al., 2015; Lun et al., 2015; Olstad et al., 2019; Fame and Lehtinen, 2020; Duy et al., 2022; Gerstmann et al., 2022; Kato and Shindo, 2024). Previous studies performed in early chicken embryos revealed that artificial leakage of CSF from the ventricles resulted in drops in intraventricular CSF pressure (P_CSF_) and abnormal morphologies of brain vesicles, such as ventricular collapses and wrinkled brain walls (Pexieder and Jelínek, 1970; Desmond and Jacobson, 1977), indicating the contribution of CSF to P_CSF_ and the necessity of P_CSF_ in the initial step of brain development. The outward or dorsal growth of the forebrain or midbrain walls suggests that they may be passively ballooned through a pressure difference across walls (i.e., higher pressure in the cavity) (Desmond and Jacobson, 1977; Garcia et al., 2019), as observed in the development of the lung (Nelson et al., 2017). Nevertheless, how P_CSF_ is created in the embryonic head and whether it varies in a stage-dependent or species-specific manner are not well understood. A recent study in mice revealed that during the period before osteogenesis begins in the skull vault, the connective and scalp tissues consisting of the epidermis and the underlying mesenchymal layer exhibit elastic and contractile properties to confine brain vesicles inwardly (Tsujikawa et al., 2022; reviewed in Miyata, 2023). Therefore, we first sought to determine the relative contributions of internal factors (i.e., CSF volume) to P_CSF_ in embryonic mice isolated from the amniotic cavity (referred to as P_CSF-ISO_) and then to determine the relative contributions of external factors (i.e., brain wall volume and confinement by the meninges or more external connective and scalp tissues) to P_CSF-ISO_. In chicken embryos, pressure in the amniotic cavity containing amniotic fluid (referred to as P_AF_) has been shown to be lower than that in P_CSF_ (Jelínek and Pexieder, 1970). Accordingly, we further sought to elucidate the relationships among P_CSF-ISO_, the intraamniotic pressure generated under the contractility of the uterus (referred to as P_AF-IU_), and the P_CSF_ of embryonic mice *in utero* (referred to as P_CSF-IU_).

Our results suggest that the contractile connective tissues surrounding the brain vesicle contribute to increasing P_CSF_ in both mouse and chicken embryos. In mice, the susceptibility of P_CSF_ to external factors was much greater because of its relationship with P_AF-IU_. At E13, P_AF-IU_ was twelve times higher than P_CSF-ISO_, and direct measurement of P_CSF-IU_ was found to be a sum of P_CSF-ISO_ and P_AF-IU_. At other stages, P_CSF-IU_ could not be directly measured. However, a constant decline in P_AF-IU_, which was much greater than P_CSF-ISO_ throughout the stages examined, suggested that P_CSF-IU_, estimated at all stages based on P_CSF-IU_ = P_CSF-ISO_ + P_AF-IU_ obtained at E13, may also decline from approximately 2,000 Pa at E10 to approximately 500 Pa at E16.

## Results and Discussion

### Stage-dependent changes in embryonic mouse P_CSF-ISO_

First, we measured P_CSF_ in mouse embryos isolated from the uterus (P_CSF-ISO_). We used a water manometer (Fig. 1A-B) modified from previously established methods (Landis 1926; Jelínek and Pexieder, 1970). Because of its structural simplicity, which requires no amplifiers or other equipment, accuracy can be reasonably expected in the measurement of P_CSF_ by the Landis micromanometer (Landis, 1926) as long as the colored saline, contained in a glass capillary, to physically face CSF is clearly visible in the embryo’s ventricle through the superficial tissue layers (i.e., brain wall and connective and scalp tissues) (Jelínek and Pexieder, 1970). The P_CSF-ISO_ values obtained (Fig. 1C) were 16.6±8.3 Pa at E10 (n=8), 38±15 Pa at E11 (n=7), 60.7±14 Pa at E12 (n=33), 82.7±18.2 Pa at E13 (n=29), 99.5±26.4 Pa at E14 (n=19), 109.4±15.5 Pa at E15 (n=9), and 106.7±8.5 Pa at E16 (n=9). The gradually increasing slope of P_CSF-ISO_ obtained in this study between E10 and E15 was consistent with those reported in previous studies of perinatal and early postnatal rodents (Jones and Bucknall, 1987).

**Figure 1.**
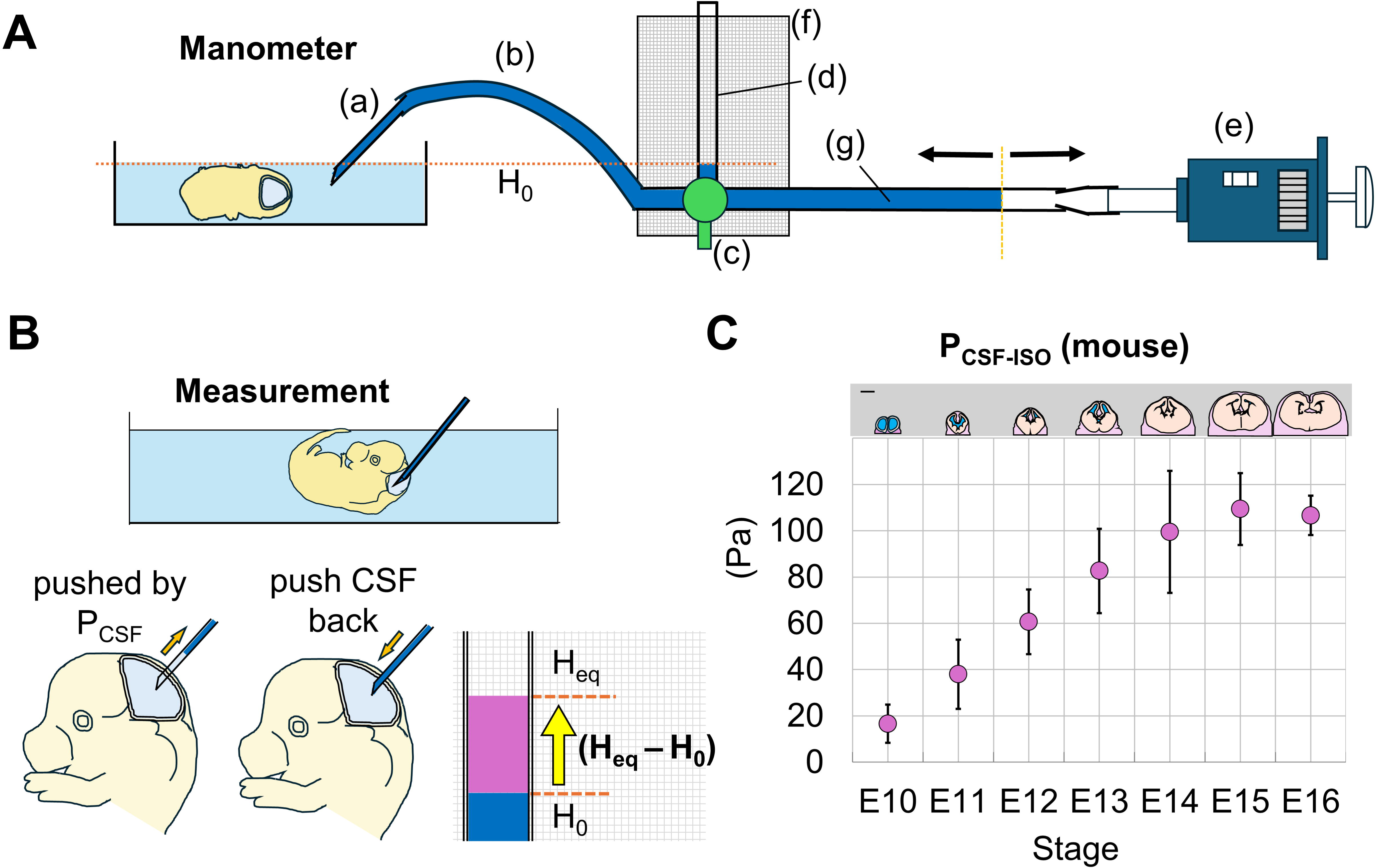
The Landis water manometer system and the obtained P_CSF-ISO_ values in mouse embryos. A. The water manometer was composed of a glass capillary needle (a), a pressure-resistant polyurethane tube (b), a three-way stopcock (c), a pressure gauge (d, outer cylinder of a 1 mL syringe), a micropipette or syringe (e), and 1 mm grid paper (f). The needles, tubes, and syringes were filled with colored saline (g). After an embryo to be measured was held onto a rubber-bottomed dish and submerged with saline (1.5 cm deep), calibration to obtain H_0_ (starting hydrostatic pressure) was performed near the head of the embryo by setting the interface between the saline in the dish and the colored saline in the capillary to the needle tip. B. Measurement of P_CSF-ISO_. Upon introduction of the needle into the midbrain ventricle, the colored saline was pushed back by CSF pressure from the tip of the needle into the proximal lumen. The pressure in the measuring system was subsequently increased until equilibrium was reached (by pushing the colored saline to the needle tip forward) to obtain H_eq_. C. P_CSF-ISO_ of mouse embryos from E10 to E16. Scale, 1mm.

### Contribution of CSF volume to embryonic mouse P_CSF-ISO_

To evaluate the contribution of CSF volume to P_CSF-ISO_, P_CSF-ISO_ measurement at E13 was coupled with intraventricular injection of saline (Fig. 2A). Injection of 1 μl did not lead to significant P_CSU-ISO_ changes (88.6±5.5 Pa [n=9, p=0.13, Welch’s t test]). Injections of 2 μl, 3 μl, 4 μl, and 5 μl resulted in significant P_CSF-ISO_ elevations (101±8.1 Pa by 2 μl [n=6, p=1.2×10^-3^, Welch’s t test], 123.9±11.4 Pa by 3 μl [n=6, p=1.9×10^-5^, Welch’s t test], 114±14.7 Pa by 4 μl [n=6, p=1.4×10^-3^, Welch’s t test], and 133.7±15.3 Pa by 5 μl [n=12, p=1.7×10^-9^, Welch’s t test]), with dose dependent increases between 2 μl and 5 μl (p=2.0×10^-5^, Welch’s t test), 3 μl and 5 μl (p=0.15, Welch’s t test) or 4 μl and 5 μl (p=0.025, Welch’s t test) (Fig. 2B). The total volume of the ventricular space in the E13 head (from the lateral ventricles to the caudal border of the fourth ventricle) was estimated based on 3D reconstructions of vibratome sections (Fig. S1A) to be approximately 3 μl (3.0±0.5 μl, n=11) (Fig. S1B), consistent with a previous measurement (Johansson et al., 2006). The fact that P_CSF-ISO_ did not so sensitively respond to a considerable level (∼30% of the original ventricular volume) of CSF increase could be explained by CSF flows to the subarachnoidal space (Akai et al., 2018), spinal cord, or extra-nervous tissues (Akai et al., 2021) and by elasticity of the scalp (Tsujikawa et al., 2022).

**Figure 2.**
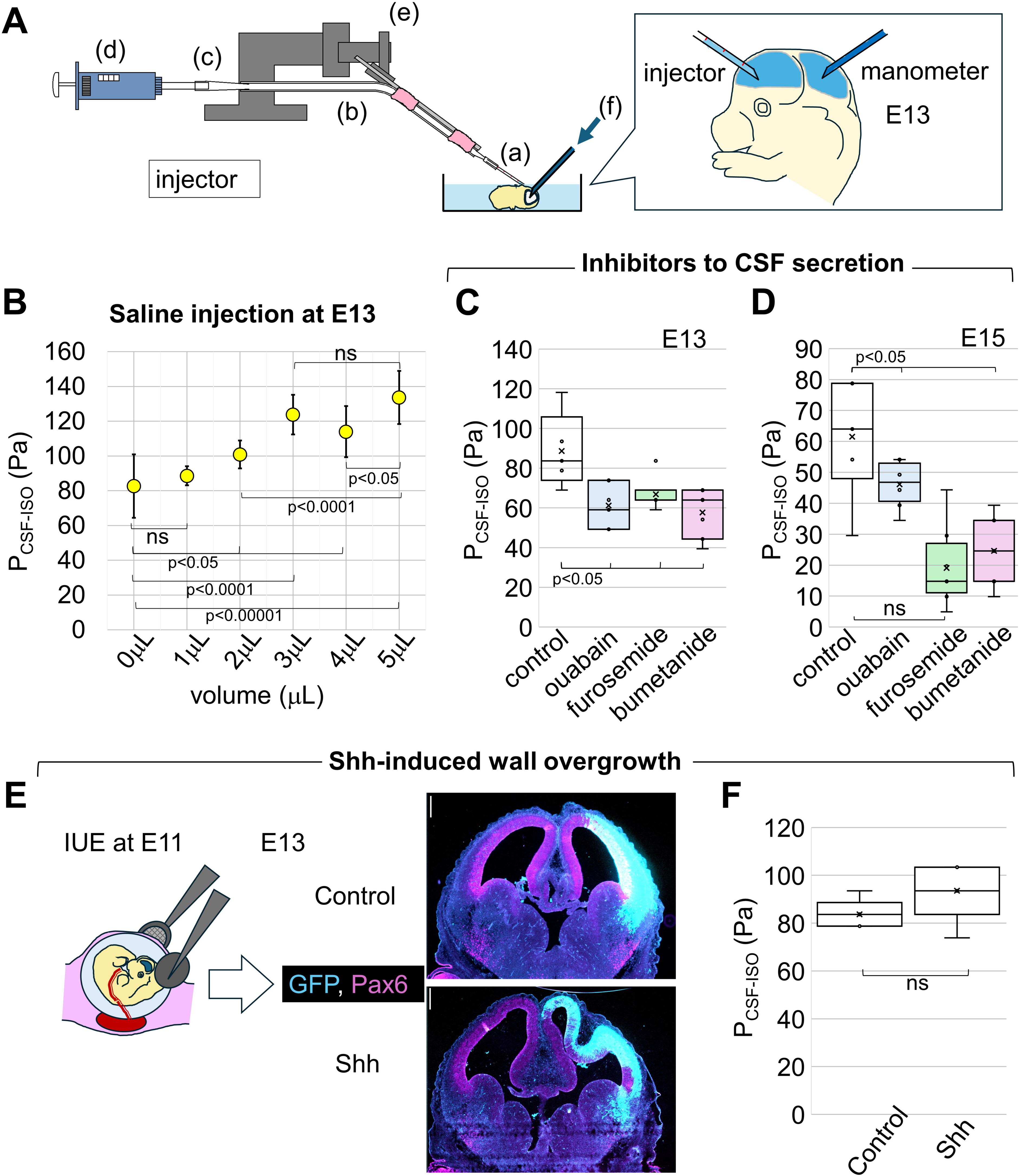
Influences of changes in the volume of CSF and brain walls on P_CSF-ISO_. A. Setup for measuring P_CSF-ISO_ (in the midbrain ventricle) while injecting saline or inhibitors (into the telencephalic [lateral] ventricle). B. P_CSF-ISO_ at E13, after intraventricular injection of saline. C. P_CSF-ISO_ at E13, after intraventricular injection of inhibitors of CSF secretion. D. P_CSF-ISO_ at E15, after intraventricular injection of inhibitors of CSF secretion E. Photomicrographs comparing coronally sectioned E13 cerebral walls between groups in which control or shh plasmid vectors were introduced at E11 via *in utero* electroporation. Scale, 0.25 mm. F. Graph comparing the dorsoventral length of the cerebral wall between the control and Shh groups at E13. G. Graph comparing the total Pax6^+^ area in the cerebral wall between the control and Shh groups at E13. H. Graph comparing the ventricular size between the control and Shh groups at E13. I. Graph comparing P_CSF-ISO_ between the control and Shh groups at E13.

We then tried to decrease the volume of CSF. Since removal of CSF using the capillary system used for saline injection was not technically possible, we took a pharmacological approach. Inhibitors to ion cotransporters or Na^+^/K^+^ ATPase have been widely used for reducing CSF secretion (Praetorius and Damkier, 2017), and the expression of the Na^+^/K^+^ ATPase (Johansson et al,, 2008) (to be blocked by ouabain), Na^+^/2Cl^−^/K^+^ cotransporters (NKCC) (Li et al., 2002) (to be blocked by furosemide), and K^+^/Cl^−^ cotransporters (KCC) (Li et al., 2002) (to be blocked by bumetanide) is reported (in the choroid plexus and neuroepithelia) already during the mid-embryonic stage that we focused. At E13, we injected 1 μl, a volume that by itself would not directly influence P_CSF-ISO_, of ouabain, furosemide, or bumetanide into the latera;/cerebral ventricle. P_CSF-ISO_ measured 1 hr later declined by ∼30% in all treatments (88.6±18.7 Pa by saline [n=5], 61.2±10.2 Pa by ouabain [n=7, p=0.02, Welch’s t test], 66.8±8 Pa by furosemide [n=7, p=0.02, Wilcoxon rank sum test], 57.7±12 Pa by bumetanide [n=7, p=0.02, Welch’s t test]) (Fig. 2C). At E15, with injection of 1.5 μl inhibitor solutions, similar reductions in P_CSF-ISO_ were observed (61.5±18.4 Pa by saline [n=6], 36.1±4.7 Pa by ouabain [n=8, p=1.2×10^-3^, Welch’s t test], 46.1±6.9 Pa by furosemide [n=8, p=0.1, Welch’s t test], 25.8±8.4 Pa by bumetanide [n=8, p=2.3×10^-3^, Welch’s t test]) (Fig. 2D). Thus, CSF volume is a contributing factor to P_CSF-ISO_.

### Influence of the cerebral wall volume increase on embryonic mouse P_CSF-ISO_

To determine whether pathological expansion and/or thickening of the brain wall and the resulting narrowing of the ventricles influence P_CSF-ISO_, we artificially induced the overproliferation of undifferentiated neural progenitor cells (NPCs) via forced expression of sonic hedgehog (shh) (Shikata et al., 2011). Coronal sections of E13 cerebral walls that had been subjected to *in utero* electroporation (IUE) with a plasmid vector to express shh at E11 demonstrated that they were dorsoventrally expanded (Fig. 2E): the length between the dorsal and ventral borders in the dorsolateral cerebral wall was greater (977.3±80.4 μm in the control [n=13], 1431.5±338 μm in shh [n=18, p=9.7×10^-9^, Wilcoxon rank sum exact test]) (Fig. S2A), and the area occupied by Pax6^+^ nuclei of NPCs was also greater (8.7×10^5^ μm^2^ in the control [n=19], 1.6×10^6^ μm^2^ in shh [n=26, p=3.3×10^-12^, Wilcoxon rank sum exact test]) (Fig. S2B).

Many ectopic (far basal) Pax6^+^ nuclei and nonsurface Pax6^+^pH3^+^ dividing cells were found in highly buckled regions, in which the most extensive overproliferation occurred per given space. Due to severe buckling of the wall, the ventricle (% area of the transfected side/area of the opposite side]) in the shh-treated group was narrowed (93±12.4% in the control [n=23], 75.7±17.7% in the shh-treated group [n=23, p=4.6×10^-4^; Welch’s t test]) (Fig. S2C). P_CSF-ISO_ measured in shh-overexpressing (overproliferative) cerebral vesicles tended to slightly increase (93.5±12.1 Pa [n=5] compared with 83.7±6 Pa [n=5, control IUE]), but this increase was not statistically significant (p=0.15, Welch’s t test) (Fig. 2F).

### Contribution of the scalp to P_CSF-ISO_ in embryonic mice and P_CSF_ in embryonic chickens

In a previous study, we reported that approximately 93 Pa was sufficient to balloon the collapsed/shrunken scalp, which was induced in E13 mouse heads by removing (aspirating) the brain to the original contour (Tsujikawa et al., 2022). Given that this value is comparable to that of P_CSF-ISO_ at E13 (82.7±18.2 Pa [Fig. 1C]), we investigated the contribution of the scalp (i.e., epidermal and mesenchymal tissues, not including pial and arachnoidal cells differentiating on the outer surface of the brain vesicle [DeSisto et al., 2020]), which is elastic and contractile (Tsujikawa et al., 2022), to P_CSF-ISO_. When the scalp was surgically removed (Fig. 3A), P_CSF-ISO_ strikingly decreased to 10.4±5.6 Pa (n=20, p=3.5×10^-9^, Wilcoxon rank sum test) (Fig. 3B) at E13. Declines in P_CSF-ISO_ by scalp removal were reproduced at E12 (from 60.7±14 Pa to 10.1±4.9 Pa [n=9, p=2.5×10^-19^, Welch’s t test]) (Fig. 3B). Separate pharmacological tests to weaken (by blebbistatin) or strengthen (by calyculin A) actomyosin-dependent contractility of the scalp, which can compressively influence the morphology of the brain (Tsujikawa et al., 2022), decreased (from 58.1±11.8 Pa [n=10] to 28.4±4.8 Pa [n=9, p=9.3×10^-6^, Welch’s t test]) or increased (from 55.6±11.1 Pa [n=10] to 83.1±15.5 Pa [n=9, p=6.0×10^-4^, Welch’s t test]) P_CSF-ISO_, respectively (Fig. 3C), at E12. At E14 and E15, scalp removal decreased P_CSF-ISO_ to 13.1±4.0 Pa at E14 [n=9, p=1.2×10^-11^, Welch’s t test] and to 36.1±11.5 Pa at E15 [n=7, p=1.0×10^-7^, Welch’s t test] (Fig. 3B). Incisions (along the anteriorDposterior axis) to the meninges (Fig. 3A) to release possible tangential tension along it did not change P_CSF-ISO_ at E13 and E14. Only at E15 was a slight decline in P_CSF-ISO_ observed (Fig. 3A). These results indicate that elastic and contractile scalp tissue plays a role, via its inward confinement of the brain, in creating P_CSF-ISO_.

**Figure 3.**
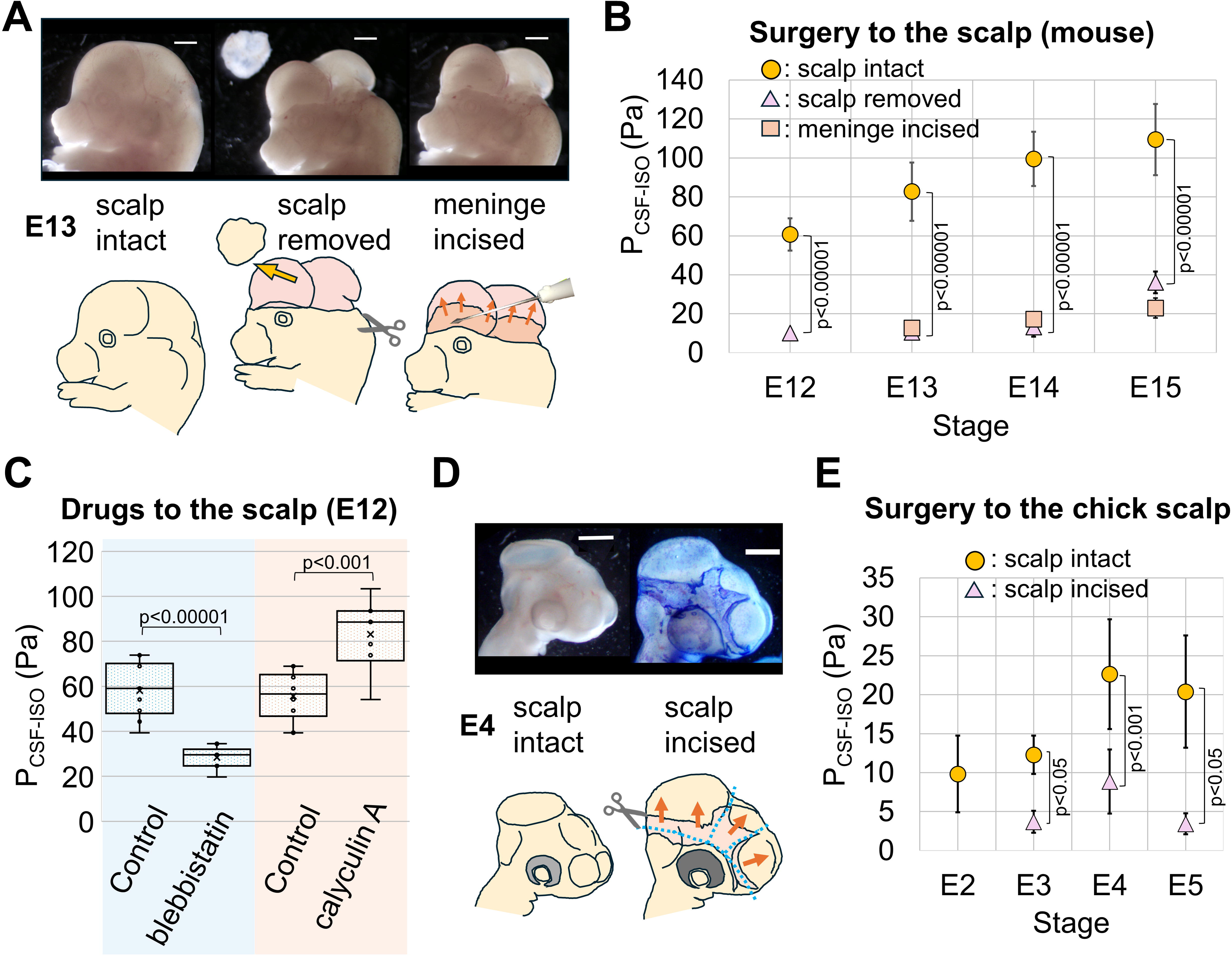
Contribution of the contractile scalp to P_CSF-ISO_. A. An E13 mouse embryo was subjected to surgery to remove the brain-surrounding scalp tissues (center) and to further incise the exposed meninges (right). Scale, 1mm. B. Graph comparing P_CSF-ISO_ between the scalp-intact, the scalp-removed and the meninges-incised groups at E12, E13, E14, and E15. C. Graph comparing P_CSF-ISO_ between the control, the actomyosin-inhibited, and the actomyosin-activated groups at E12. D. The brain-surrounding scalp tissues of E4 chicken embryos were incised via surgery. Photomicrograph of an E4 chicken embryo after incision, toluidine blue-stained, showing a wound where the brain was exposed. Scale, 1mm. E. Graph comparing P_CSF_ in chicken embryos between the scalp-intact and the scalp-incised groups at E2, E3, E4, and E5.

We further investigated whether P_CSF_ is also significantly influenced by the scalp in E3-E5 chicken embryos. P_CSF_ values measured in the scalp intact chicken embryos (9.8±4.9 Pa at E2 [n=3], 12.3±2.5 Pa at E3 [n=7], 22.6±7 Pa at E4 [n=10], and 20.4±7.2 Pa at E5 [n=7]) (Fig. 3D-E) were consistent with values reported in previous studies (Jelínek and Pexieder, 1970; Garcia et al., 2018). When incisions were carefully made onto the embryonic chicken scalp tissues outside the brain wall (only to the right side that was accessible) (Fig. 3D), the wound immediately started to open perpendicularly to the incision, suggesting a preexisting tensile (prestretched) property. Reflecting the loss of such tangential tension upon incisions to the scalp, P_CSF_ dropped to 20-40% of the original level (to 3.7±1.4 Pa [n=4, p=0.009, Wilcoxon rank sum test] at 3 d, to 8.9±4.1 Pa [n=5, p=4.6×10^-4^, Welch’s t test] at 4 d, and to 3.4±1.3 Pa [n=5, p=4.5×10^-3^, Wilcoxon rank sum test] at 5 d) (Fig. 3E). Thus, the contribution of brain-surrounding, contractile scalp tissue to the creation of P_CSF_ is common in mammals and birds during development.

### Measurement of P_AF-IU_

Given the abovementioned susceptibility of P_CSF-ISO_ to the externally provided physical load, we sought to determine whether and, if so, how the intraamniotic pressure generated under the contractility of the uterus (referred to as P_AF-IU_) affects P_CSF_. To first measure P_AF-IU_, we used two different methods (Fig. 4A): a water manometer was used from E10 to E18, whereas a pressure sensor (transducer) was used from E13 to E18. The obtained P_AF-IU_ values (Fig. 4B) were: 2,024.6±211.6 Pa at E10 (n=7), 1,701.2±179.6 Pa at E11 (n=10), 1,312.7±90.6 Pa at E12 (n=6), 1,030.7±257.5 Pa by manometer (n=8) and 1,277.7±286.5 Pa by sensor (n=10) at E13, 698.2±205.8 Pa by manometer (n=12) and 1,015.8±144.6 Pa by sensor (n=7) at E14, 676±64.8 Pa by manometer (n=9) and 653.2±121.7 Pa by sensor (n=6) at E15, 448±85 Pa by manometer (n=6) and 486.2±190.8 Pa by sensor (n=10) at E16, 434.8±26.2 Pa by manometer (n=6) and 397.2±91.2 Pa by sensor (n=8) at E17, and 222.2±116.1 Pa by manometer (n=7) and 495±125.3 Pa by sensor (n=7) at E18. Despite slight differences between the manometer-measured P_AF-IU_ and the sensor-measured P_AF-IU_ at a few embryonic days, both methods consistently showed a stage-dependent decline in the P_AF-IU_ from 2 kPa to 0.5 kPa (from E10 to E16). The individual values of P_AF-IU_ (except at E10) and the slopes formed by these values (from E11 to E15) are very similar to the results shown previously in embryonic mice (McCafferty et al., 1964).

**Figure 4.**
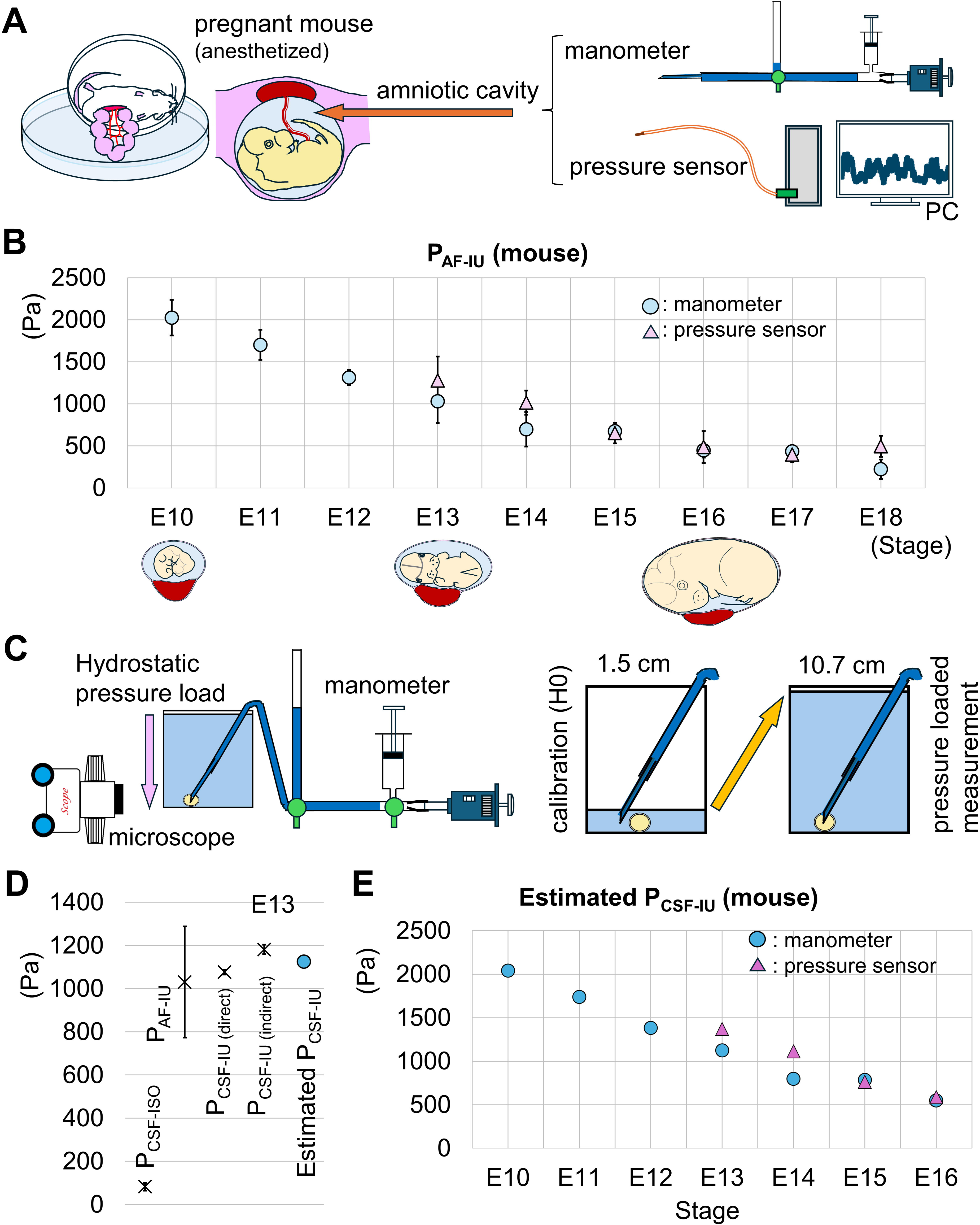
Stage-dependent changes in P_AF-IU_ and its contribution to P_CSF-IU_. A. Setup for measuring P_AF-IF_. B. Graph depicting P_AF-IU_ measured via a water manometer (E10∼E18) or a pressure sensor (E13∼E18). C. Measurement of P_CSF_ in isolated E13 embryos under loading of hydrostatic pressure corresponding to P_AF-IU_. After calibration at 1.5 cm, the embryo was deeply (10.7 cm) submerged in saline. D. Graph comparing P_CSF-ISO_ at E13, P_AF-IU_ at E13, P_CSF_ directly measured *in utero* at E13, and P_CSF_ measured in isolated E13 embryos under pressure loading corresponding to E13 P_AF-IU_, and P_CSF-IU_ estimated from P_CSF-ISO_ + P_AF-IU_ at E13. E. Graph showing the stage-dependent changes in the estimated P_CSF-IU_ (= P_CSF-ISO_ + P_AF-IU_).

Throughout our examinations, P_AU-IF_ was much higher than P_CSF-ISO_. The earlier the stage, the greater the P_AU-IF_ dominance over P_CSF-ISO_: 100-times at E10, 43-times at E11, 23-times at E12, 12-times at E13, six-times at E14, and five-times at E15 and E16.

### Direct measurement of P_CSF-IU_ at E13

We hypothesized that such a relatively high P_AF-IU_ acts physically as a strong inducer of P_CSF-IU_, as contraction of the cardiac or smooth muscles elevates vascular pressure; therefore, we sought to determine whether there is a P_CSF-IU_ = P_CSF-ISO_ + P_AF-IU_ relationship. The P_CSF-IU_ measured via our manometer was 1,076.4±11.4 Pa (n=3), which was slightly lower than the value expected (1,113.4 Pa) by simply summing 82.7 Pa (P_CSF-ISO_) and 1,030.7 Pa (P_AF-IU_). A small opening (1.5-2 mm in diameter) was made in the uterine wall to secure the visibility needed for the manometer (to monitor the colored saline in the ventricle of the embryo through the amniotic membrane), which could have slightly weakened the overall confinement by the entire uterine tube. Taking that possibility into account, this result of direct measurement at the E13 is consistent with our model of P_CSF-IU_ = P_CSF-ISO_ + P_AF-IU_.

### PCSF measurement in isolated E13 mice under experimental hydrostatic pressure loading

To further examine the physical susceptibility of P_CSF_ to external factors and evaluate the aforementioned model for pressure summation, we next took a different approach. Isolated E13 mouse embryos were subjected to P_CSF_ measurement under externally loaded hydrostatic pressure corresponding to E13 P_AF-IU_. After calibration at the height of an E13 embryo held on a rubber plate attached to the bottom of a water chamber, a 10.7 cm depth of saline was provided to that embryo (Fig. 4C). With this pressure-loading device mimicking the intraamniotic (*in utero*) space, the P_CSF_ was 1,181.4±22 Pa (n=6), comparable with the directly measured P_CSF-IU_. This result confirmed the physical susceptibility of P_CSF_ to P_AF-IU_, again supporting our model for pressure addition: P_CSF-IU_ = P_CSF-ISO_ + P_AF-IU_ (Fig. 4D).

## Conclusions

Using a Landis manometer, we first measured the intraventricular CSF pressure in mouse embryos isolated from the amniotic cavity (P_CSF-ISO_). We found contributions of the CSF volume and the confinement from the brain-surrounding connective/scalp tissues to P_CSF-ISO_ (Fig. 5).

**Figure 5.**
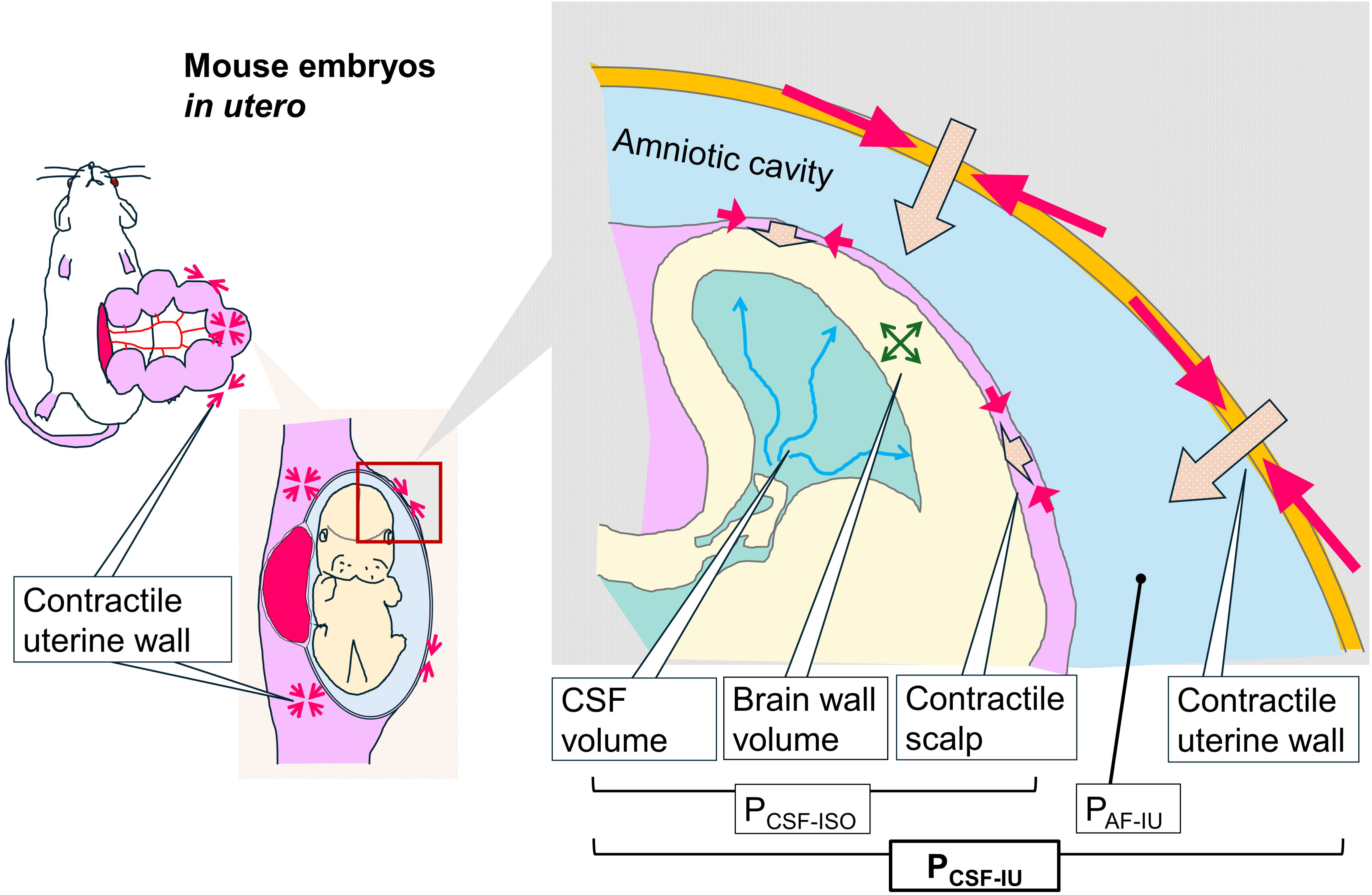
The contributions of the liquid volume and the surrounding tissues to pressure. The CSF volume and the contractile of the scalp generates P_CSF-ISO_, and the contractile of the uterine wall generates P_AF-IU_. With these physical influences, the interventricular space of mouse embryos in utero are highly pressurized (P_CSF-IU_).

Further analysis on the CSF pressure in mouse embryos *in utero* (P_CSF-IU_) and the amniotic pressure *in utero* (P_AF-IU_) revealed that mouse embryos *in utero* are growing under a significant physical influence from the contractile uterine wall, with their intraventricular space highly pressurized (Fig. 5).

## Limitations and perspectives

At this point, it is uncertain whether the rule of pressure addition (P_CSF-IU_ = P_CSF-ISO_ + P_AF-IU_) found at E13 can be applied to other embryonic ages. For example, ossification in the skull vault, which occurs during late embryonic days (Rice et al., 2000), could buffer the potential influence of P_AF-IU_ on P_CSF-IU_ across brain-surrounding head tissues. Nevertheless, the almost constant decline in P_AU-IU_ and the consistent (although shrinking) dominance of P_AU-IF_ over P_CSF-ISO_ (100-times greater at E10 ➔ five-times greater at E15 and E16) strongly suggest that P_CSF-IU_ may be greater in early embryonic days than in mid- or late embryonic days in mice. P_CSF-IU_ in embryos can be estimated as follows: 2,020 Pa at E10, 1,750 Pa at E11, 1,400 Pa at E12, 1,100 Pa at E13, 800 Pa at E14, 800 Pa at E15, and 500 Pa at E16 (Fig. 4E). High P_CSF-IU_ in early embryonic brain vesicles caused by high P_AF-IU_ is mechanically reasonable because it provides a hydrostatic skeleton to a thin and fragile wall of the brain vesicle (Kier, 2012; Tsujikawa et al., 2022; Adameyko, 2023). It will be interesting to explore how this apparently high P_CSF-IU_ during the early brain-forming stage could be associated with active proliferation and/or differentiation through the functions of mechanosensory proteins such as Piezo1 (Pathak et al., 2014; Gudipaty et al., 2017; He et al., 2018; Nourse et al., 2022) and mechanochemical coupling (Hannezo and Hesenberg, 2019).

## Materials and Methods

### Animals

The animal experiments were conducted according to the Japanese Act on Welfare and Management of Animals, the Guidelines for Proper Conduct of Animal Experiments (published by the Science Council of Japan), and the Fundamental Guidelines for Proper Conduct of Animal Experiments and Related Activities in Academic Research Institutions (published by the Ministry of Education, Culture, Sports, Science and Technology, Japan). All protocols for the animal experiments were approved by the Animal Care and Use Committee of Nagoya University (No. 29006). Pregnant female mice (*Mus musculus*) were obtained from SLC (Hamamatsu, Japan; for ICR mice) or by mating at Nagoya University. Embryonic day (E) zero was defined as the day of vaginal plug identification. Fertilized chicken (*Gallus gallus domesticus*, Hypeco near) eggs were obtained from the Yamagishi poultry farm (Mie, Japan) and incubated at 38.5 °C.

### Measurement of the intraventricular pressure (P_CSF_) in mouse or chick embryos

#### Micromanometer

CSF pressure in the ventricular space of embryonic brain vesicles was measured using a water micromanometer according to previously established methods (Landis 1926; Jelínek and Pexieder, 1970). Briefly, a pressure-resistant polyurethane tube (∼217 kPa, inner diameter 3 mm, outer diameter 5 mm: Saint-Gobain, Japan) with an adapter (pipette tip and silicone rubber tube; inner diameter 1 mm, outer diameter 3 mm: Monotaro, Japan) at one end to attach a glass capillary needle (outer diameter 1 mm, GD-1, Narishige, Japan; pulled with a Narishige PN-30 tip puller and polished with a Narishige EG-400 micro grinder to an outer diameter of 300-500 μm with a 30-degree angle at the tip) to be inserted into the ventricle was connected to the pipette tip of a Pipetman P1000 pipette (Gilson, WI, US.) via a three-way stopcock (TS-TR1K, Terumo, Tokyo, Japan). While two horizontal branches of the three-way stopcock were used to send saline (Otsuka, Tokushima, Japan) containing 0.03% Fast Green FCF (Wako, Osaka, Japan) to the glass capillary, the third upright branch was used to make a pressure gauge with a 1 ml syringe (SS-01T, TERUMO, Tokyo, Japan) and a 1 mm grid paper (KOKUYO, Tokyo, Japan) (Fig. 1A).

#### P_CSF_ measurement procedures

Mouse (E10-E16) embryos were isolated from the uterus and kept in a Petri dish (10-15 cm) containing DMEM/Ham’s F-12 culture medium (Fujifilm Wako, Osaka, Japan). Each embryo was transferred to another medium-containing Petri dish (10 cm) and gently held and immobilized on a silicone rubber plate (∼ 5 mm thick, made over the bottom surface of the Petri dish by mixing KE-103 and CAT-103 from Shin-Etsu silicones [Shin-Etsu Chemical, Tokyo, Japan]) via stainless insect pins (without piercing through or pressing the embryo). The depth (8∼10 mm) of the medium was set to soak the entire mouse embryo (see below). For *in ovo* measurements of chicken (E2∼E5) embryos, the chorion and amnion were carefully removed from above the head. P_CSF_ was measured in living embryos whose health was confirmed through observations of heart beating and/or subcutaneous blood flow. In accordance with Jelínek and Pexieder (1970), the calibration and measurement were performed as follows. At the horizontal level of each mouse or chick embryo’s head (regarded as the level of zero pressure for manometer calibration), the tip of the micromanometer (the glass capillary) was placed in the neighborhood of the future punctate of the embryo (in DMEM/F-12 [mouse] or in the amniotic fluid [chick]). Then, under a dissecting microscope (MZ7.5, Leica, Wetzlar, Germany), the pressure in the micromanometer system was calibrated by turning the thumbwheel of the pipette such that the Fast Green colored saline did not flow out of the tip of the glass capillary nor was the surrounding medium or amniotic fluid sucked into the needle. The corresponding value on the pressure gauge (the height of the colored saline) represented the starting (zero) height (H_0_). Afterward, the tip of the glass capillary needle was inserted into the midbrain vesicle. Embryos damaged during this needle insertion step were excluded from further analysis. When the needle was successfully inserted, the colored saline was automatically pushed deeper into the lumen of the glass capillary by the pressurized CSF coming from the ventricle (Fig. 1B). Then, we increased the pressure in the manometer system by turning the thumbwheel of the pipette until the CSF was pushed back from the tip of the glass capillary, and the height of the colored saline when the pressure equilibrium was thus reached (H_eq_) was read on the grid paper with a resolution of 0.5 mm (0.25 mm in chicks) (Fig. 1B). If the manometer is pressurized too quickly, the resistance of the colored saline in the system increased because of the thin tip of the glass capillary, resulting in a too-high pressure. Therefore, pressure was applied slowly. In the case of mouse embryos with scalps, measurements were taken in approximately 10 minutes.

P_CSF_ was obtained according to:

P_CSF_ = (H_eq_ – H_0_) × g × ρ (Pa)

H_eq_ – H_0_ : pressure in mmH_2_O

g: 9.803 (Pa / mmH_2_O)

ρ: 1.0043 (specific gravity of saline with 0.03% Fast Green)

When the depth of the medium surrounding an E13 mouse embryo was altered from 10 mm to 0 mm (with a water drop for obtaining H_0_ placed away from the head), the P_CSF_ did not significantly change.

### P_CSF_ measurement combined with surgeries to the scalp and the meninges

All surgical procedures for embryos were performed in culture medium (DMEM/Ham’s F-12, Fujifilm Wako, Osaka, Japan) under a dissecting microscope (Leica MZ7.5) (Fig. 3A, 3D).

Incisions in the scalps of E12∼E15 mouse embryos and E2∼E5 chicken embryos were made using microscissors (tip diameter 0.05 mm, cutting edge 3Dmm; 15D000–00, Fine Science Tools, B.C., Canada). Care was taken not to damage the meninges or brain. When the scalp was incised along the anteriorDposterior axis, the cut edges of the scalp were immediately separated along the dorsoventral axis of the head as previously described (Tsujikawa et al., 2022). The circularly incised scalp was then removed from each embryo. To make bilateral incisions to the meninges after scalp removal, the meninges covering the lateral part of the cerebrum and midbrain were incised in an anterior-to-posterior direction using an ophthalmic scalpel (8065912001, 20G Size, Alcon, Geneva, Switzerland). Comparisons were made between P_CSF_ values measured in untreated control embryos and P_CSF_ values measured at 20∼30 min in these surgically treated embryos.

### P_CSF_ measurement combined with intraventricular fluid injection

The intravenous injection of DMEM/Ham’s F-12 culture medium was performed via a manual injection system as follows. A pipette tip attached to a Pipetman P1000 was connected through a pressure-resistant polyurethane tube (∼1034 kPa, inner diameter 2 mm, outer diameter 4 mm: Monotaro, Japan) to a pulled glass capillary (GD-1, Narishige, Tokyo, Japan) via an adapter (pipette tip and silicone rubber tube; inner diameter 1 mm, outer diameter 3 mm: Monotaro, Japan) (Fig. 3A). The body (nontapered part) of the glass capillary was marked to indicate steps of 1 μl volume (every 3.5 mm). The tube was bundled to a micropipette injection holder (HI-7, Narishige, Tokyo, Japan) with waterproof tape (Nichiban, Tokyo, Japan) and attached to a manual manipulator (MWS-2, Narishige, Tokyo, Japan). The positions of the needles were optimized using a manipulator. After each mouse embryo was gently immobilized on a silicone rubber plate, medium (1-5 μl) was injected into the cerebral vesicle (lateral ventricle) by turning the thumbwheel of the micropipette. Soon (within 5 min) after injection, P_CSF_ in the midbrain ventricle was measured via a water micromanometer.

### P_CSF_ measurement combined with pharmacological examinations

E12 mouse embryos were incubated for 60 minutes in culture medium containing 0.5% DMSO (SigmaDAldrich, MO, US.) only, 0.5% DMSO plus 100 μM blebbistatin (Calbiochem, CA, US.), or 100 nM calyculin A (Fujifilm Wako, Osaka, Japan), after which P_CSF_ was measured.

Throughout the incubation of the embryos, the medium was carefully oxygenated with 95% O_2_/5% CO_2_ to prevent the intramesenchymal or perimeningeal vessels from contracting in response to hypoxia. To examine the effects of inhibitors of CSF secretion on P_CSF_, 5 mM furosemide (ab120314, Abcam, Cambridge, UK), 1 mM bumetanide (ab142489, Abcam, UK), 1 mM ouabain (ab120748, Abcam, UK), or saline was injected (1 μl at E13, 1.5 μl at E15) into the lateral ventricle in utero, and P_CSF_ was measured 60 min later.

### Measurement of the intraamniotic fluid pressure *in utero* (P_AF-IU_)

#### P_AF-IU_ measurement preparation

A pregnant mouse (E10∼E18) was anesthetized intraperitoneally with the mixture of medetomidine hydrochloride (0.75 mg/kg, NIPPON ZENYAKU KOGYO, Fukushima, Japan), midazolam (4 mg/kg, Sandoz, Tokyo, Japan), and butorphanol tartrate (5 mg/kg, Meiji Animal Health, Tokyo, Japan). The abdominal wall was then incised. The 10 cm dish lid was inverted and placed on a 15 cm dish filled with phosphate-buffered saline (PBS) to a depth of 10 mm. Each mouse was placed on a 10 cm lid while all the uterine horns were carefully exposed to PBS to prevent drowning during the experiment (Fig. 4A). The uterus was kept moist with PBS throughout the measurement. The mother’s heartbeat and breathing were maintained throughout.

#### P_AF-IU_ measurement using a micromanometer

The intraamniotic pressure *in utero* was measured using a water micromanometer as described for the P_CSF_ measurement. The pressure gauge was replaced by a pressure-resistant tube supported vertically by a 40 cm rod instead of a 1 mL syringe. The procedures from calibration to measurement were the same as those for the P_CSF_ measurement, but the height of the water column when colored saline was pushed with a P-1000 pipette was limited to approximately 70 mm. Therefore, a three-way stopcock was attached between the pipette tip and tube, and a 5 mL syringe was attached to the branch. A glass capillary was inserted into the amniotic cavity through the uterine wall. The first P_AF-IU_ measurement was a pilot measurement, in which colored saline was slowly pushed with the syringe, and a small leakage into the amniotic cavity was confirmed with a dissection microscope. Next, in the actual measurement, colored saline was slowly pushed with the syringe to approximately 20-30 mm below this height and then slowly pushed with the pipette until the saline was pushed back to the amniotic cavity.

#### P_AF-IU_ measurement using a pressure sensor microcatheter

A microtransducer-mounted pressure sensor catheter system (signal conditioner: EVO-SD-2-FPI, sensor catheter: 0.9 Fr, FOP-LS-PT9-10, FISO Technologies, Quebec, Canada) was also used for measurement of the intraamniotic fluid pressure of the amniotic cavity *in utero.* The tip (transducer) of the sensor catheter was placed into the lumen of a 21GD23G injection needle (Terumo, Tokyo, Japan), which was inserted into the amniotic cavity. Once the catheter tip entered the amniotic cavity, the outer needle was removed. Care was taken not to remove the catheter. During measurement, the catheter was held in place to avoid contact with the embryo or amniotic membrane.

#### Data analysis of the pressure sensor microcatheter

The pressure data obtained by the sensor catheter were recorded in a CSV file on a computer via the sensor module and graphed using Microsoft Excel (Microsoft, CA, US.), and the amniotic fluid pressure was calculated by averaging the values in the flat parts of the waveform during intermittent uterine contractions.

### Direct measurement of the intraventricular pressure of embryos *in utero* (P_CSF-IU_)

The uterine horns of the anesthetized pregnant mouse were exposed on a 15 cm dish filled with PBS as in the Piu measurement procedure, and P_CSF_ was measured with a water micromanometer. The glass capillary of the water micromanometer was inserted into the midbrain vesicle through the myometrium under a dissecting microscope (Leica MZ7.5). To ensure the visibility of the colored saline, a small hole (approximately 2 mm in diameter) was made in the myometrium with microscissors (tip diameter 0.05 mm, cutting edge 3Dmm; 15D000–00, Fine Science Tools, B.C., Canada). This procedure did not change the overall morphology or physical condition of the uterine tube. After confirming the position of the brain vesicle through the exposed amniotic membrane, the glass capillary of the water micromanometer was inserted.

### Measurement of P_CSF_ in isolated embryos under pressure loading corresponding to P_AF-IU_

P_AF-IU_ was reproduced in a device to provide hydrostatic pressure (Fig. 4C). To the bottom of a tall transparent plastic chamber (7.5 cm × 7.5 cm × 12 cm), a layer of silicone rubber was made by mixing KE-103 and CAT-103 from Shin-Etsu silicones (Shin-Etsu Chemical, Tokyo, Japan). E13 mouse embryos isolated from the uterus were gently held and immobilized onto rubber plates via insect pins. The embryo was submerged in PBS at a depth of 1.5 cm. After calibration was performed (using the micropipette, by hand A) at a height of 1.5 cm near the embryo’s head, PBS was added to increase the depth to 10.7 cm. Then, under careful observation with a stereomicroscope from the sidewall of the chamber (the objective lens originally set upright was rotated 90 degrees and fixed to the chamber at a certain distance), the glass capillary needle attached to a holder was manually inserted into the midbrain vesicle of the embryo (by hand B). Then, the colored saline, which was pushed into the lumen of the manometer system to a considerable degree, was pushed back to the needle tip (using the micropipette, by hand A), followed by closure of the three-way stopcock (by hand A) and reading of H_eq_.

### 3D reconstructions of the total volume of the ventricular space

Mouse embryo at E13 was fixed by 4% PFA and the head was mounted in a block of gel containing 2.5% agarose (NIPPON GENE, Tokyo, Japan). Coronal sections 300 μm thick were prepared using a vibratome. On each slice, the front and back were photographed under a dissecting microscope. Ventricular areas were measured with ImageJ (National Institute of Health) from the lateral ventricles to the caudal border of the fourth ventricle. The terminal end was treated as a cone and the volume was determined from the area and depth of the ventricles. The middle was treated as a columnar body and the volume was determined from the average area of the front and back surfaces and the thickness of the slice (Fig. S1A).

### Immunofluorescence

Heads of embryonic mice were fixed with 4% paraformaldehyde (PFA) fixative (4 °C, 12 h). After immersion in 20% sucrose, the heads were embedded in OCT compound (Sakura Finetek, Japan), frozen and sectioned coronally (16 μm, with Leica CM1860, Wetzlar, Germany). The frozen sections were treated with the following primary antibodies: anti-GFP (chick, GFP-1020, Aves Labs); anti-Pax6 (rabbit, PRB-278O, COVANCE, 1:500); and anti-pH3 (rat, ab10543, Abcam, 1:500). After being washed, the sections were treated with Alexa Fluor 488, Alexa Fluor 546, and Alexa Fluor 647 conjugated secondary antibodies (Molecular Probes, A-10040, A-21209, A-21247, 1:1,000), and subjected to fluorescence microscopy (Olympus BX60, Tokyo, Japan).

### *In utero* electroporation

The induction of artificial overproliferation of undifferentiated NPCs was achieved by transfection with pCAGGS-ShhN-iresEGFP plasmid vector (0.5 μg/μl, a gift from Jun Motoyama [Shikata et al., 2011]) by *in utero* electroporation (IUE) of E11 mouse embryos (Fig. 2E), as described previously (Okamoto et al. 2013). After the DNA mixture was injected into the lateral ventricle, the head of the embryo in the uterus was placed between the discs of a forceps-type electrode (disc electrodes of 1 mm; CUY560P1, NEPA GENE, Chiba, Japan), and electric pulses (50 V) were applied four times, resulting in gene transfection into the cerebral wall. The pCAG-EGFP plasmid vector (0.5 μg/μl) was transfected as a control.

### Image analysis and quantification

#### Image analysis

Using ImageJ (National Institute of Health), we measured the length and area from the images taken with a microscope. In the experiment, we took a photograph of a scale, measured the intervals between the scale marks on the photograph more than 10 times, and used the average value as the standard for length.

#### Statistical analysis

The statistical analysis performed in this study compared the central tendencies of two populations based on samples of unrelated data. For this purpose, Welch’s test was prioritized (Ruxton et al., 2006) on the basis of the normal sample distribution in each group. The ShapiroDWilk normality test revealed that in most of the experimental groups in this study, the distribution was normal, and comparisons of the groups were accordingly conducted using Welch’s test. In a few cases where distribution normality was not found, the Wilcoxon rank sum exact test was used.

## Acknowledgement

We thank Takeo Matsumoto for manometer instructions, Jun Motoyama for plasmids, Namiko Noguchi and Makoto Masaoka for excellent technical assistance, and members of Miyata laboratory for discussion.

## Author contributions

K. T. and R.M. carried out experiments, analyzed the data. K.T. and T.M. designed the study and wrote the manuscript. All authors approved the manuscript.

## Research funding

JSPS 22J14017 (K.T.), JSPS 24K02205 (T.M.), MEXT 23H04306 (T.M), JSPS 23K18339 (T.M.), MEXT 21H00363 (T.M.), JSPS 21H02656 (T.M.)

---

**Figure S1.**
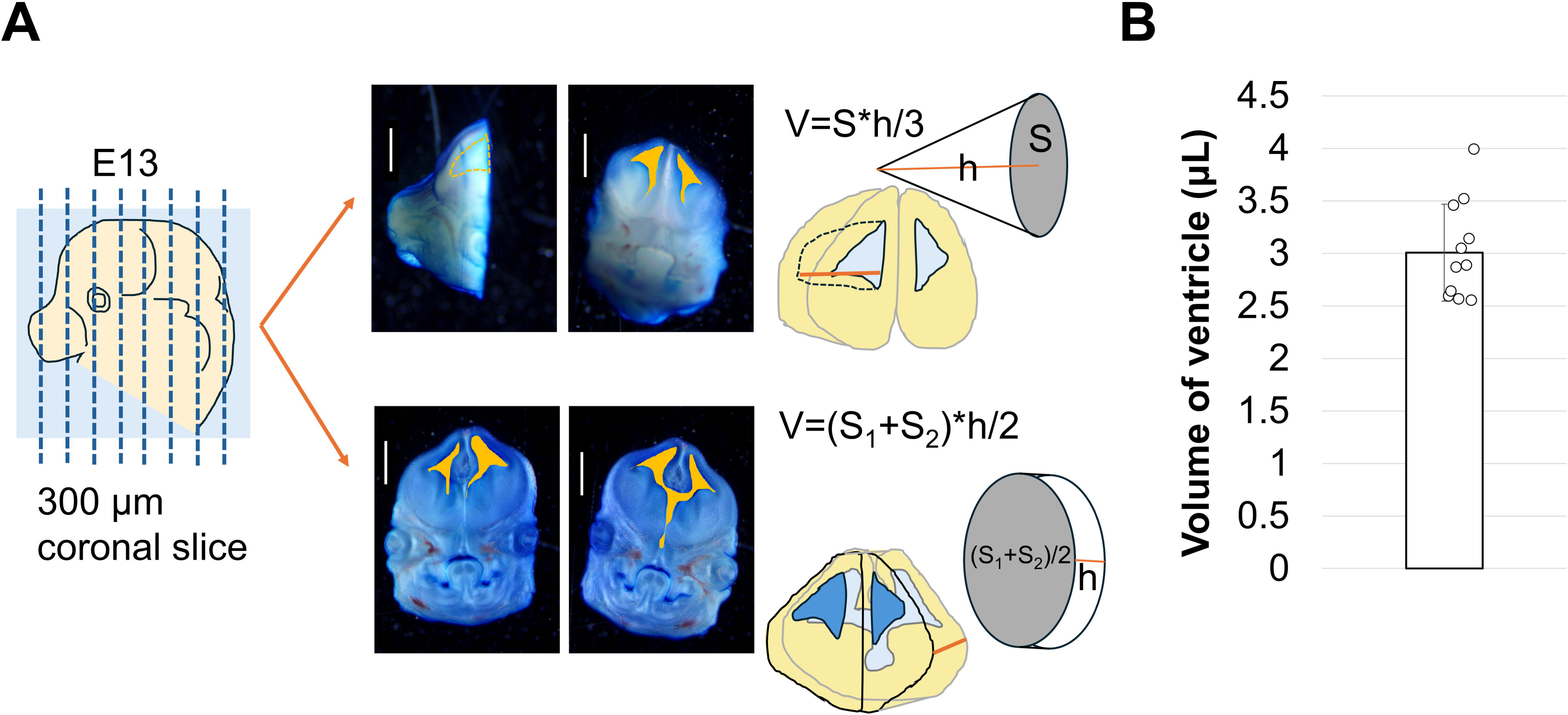
A. Ventricular volumes were measured with ventricular areas and thickness from coronal sections of mouse embryo. The terminal end was treated as a cone and the middle was treated as a columnar body. B. Graph showing the ventricular volume at E13.

**Figure S2.**
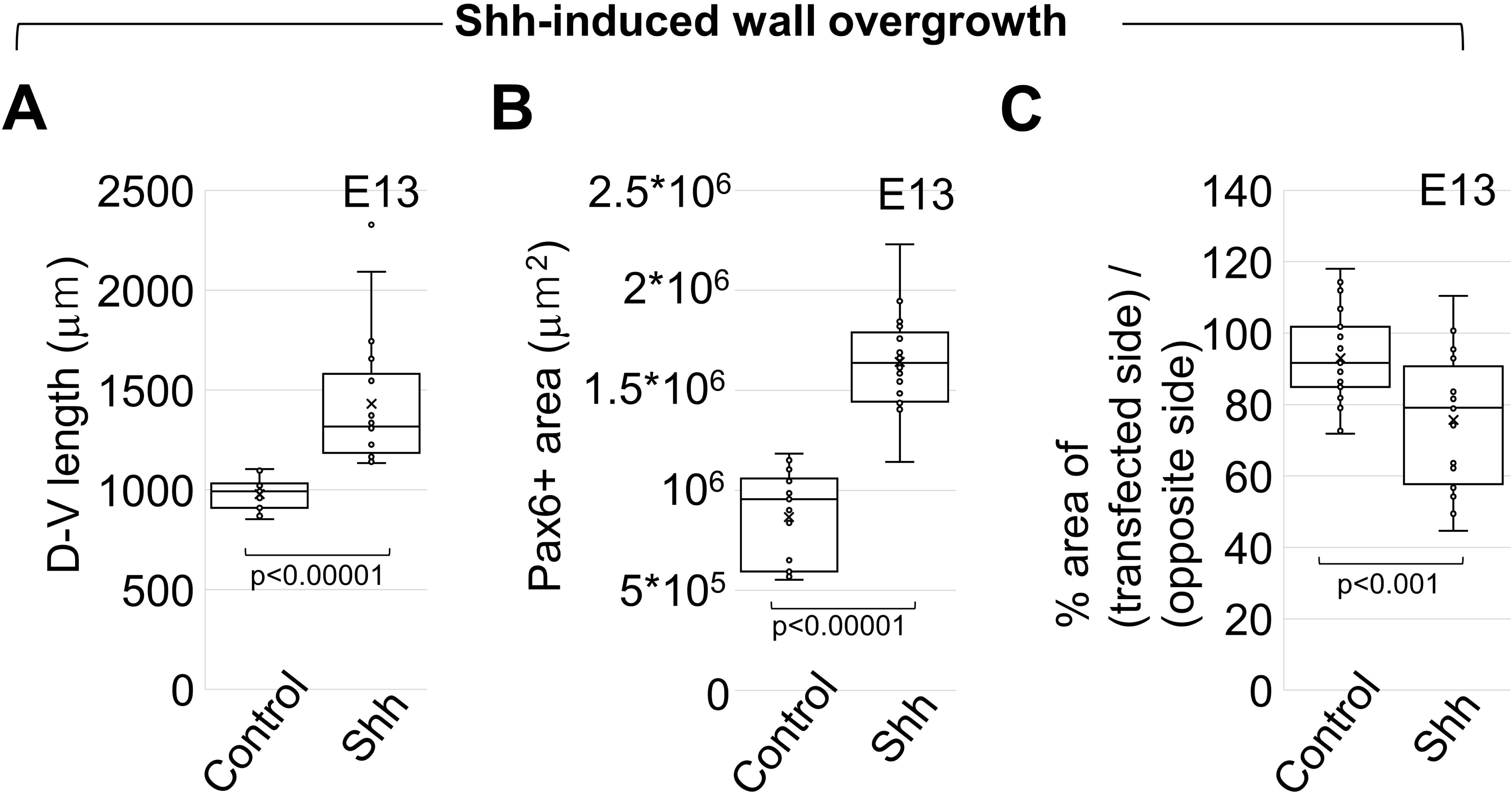
A. Graph comparing the dorsoventral length of the cerebral wall between the control and Shh groups at E13. B. Graph comparing the total Pax6+ area in the cerebral wall between the control and Shh groups at E13. C. Graph comparing the ventricular size between the control and Shh groups at E13.

## References

1. Adameyko I (2023). Evolutionary origin of the neural tube in basal deuterostomes. Curr. Biol. 33, R319–R331.

2. Akai T, Hatta T, Shimada H, Mizuki K, Kudo N, Hatta T, Otani H (2018). Extracranial outflow of particles solved in cerebrospinal fluid: Fluorescein injection study. Congenit Anom (Kyoto) 58:93–98

3. Akai T, Hatta T, Sakata-Haga H, Yamamoto S, Otani H, Yamamoto S, Kuroda S. (2021) Cerebrospinal fluid may flow out from the brain through the frontal skull base and choroid plexus: a gold colloid and cadaverine injection study in mouse fetus. Childs Nerv Syst 37:3013–3020.

4. Chau KF, Springel MW, Broadbelt KG, Park HY, Topal S, Lun MP, Mullan H, Maynard T, Steen H, LaMantia AS et al (2015) Progressive differentiation and instructive capacities of amniotic fluid and cerebrospinal fluid proteomes following neural tube closure. Dev Cell 35: 789–802

5. DeSisto J, O’Rourke R, Jones HE, Pawlikowski B, Malek AD, Bonney S, Guimiot F, Jones KL, Siegenthaler JA (2020) Single-cell transcriptomic analyses of the developing meninges reveal meningeal fibroblast diversity and function. Dev Cell 54: 43–59

6. Desmond ME, Jacobson AG (1977) Embryonic brain enlargement requires cerebrospinal fluid pressure. Dev Biol 57: 188–198

7. Duy PQ, Rakic P, Alper SL, Butler WE, Walsh CA, Sestan N, Geschwind DH, Jin SC, Kahle KT (2022). Brain ventricles as windows into brain development and disease. Neuron 110:12–15.

8. Fame RM, Lehtinen MK (2020) Emergence and developmental roles of the cerebrospinal fluid system. Dev Cell 52: 261–275

9. Fisk NM, Ronderos-Dumit D, Tannirandorn Y, Nicolini U, Talbert D, Rodeck CH. Normal amniotic pressure throughout gestation. Br J Obstet Gynaecol. 1992 Jan;99(1):18–22.

10. Garcia KE, Stewart WG, Espinosa MG, Gleghorn JP, Taber LA (2019) Molecular and mechanical signals determine morphogenesis of the cerebral hemispheres in the chicken embryo. Development 146: dev174318

11. Gato A, Desmond ME (2009) Why the embryo still matters: CSF and the neuroepithelium as interdependent regulators of embryonic brain growth, morphogenesis and histogenesis. Dev Biol 327: 263–272A

12. Gerstmann K, Kindbeiter K, Telley L, Bozon M, Reynaud F, Théoulle E, Charoy C, Jabaudon D, Moret F, Castellani V (2022) A balance of noncanonical Semaphorin signaling from the cerebrospinal fluid regulates apical cell dynamics during corticogenesis. Sci Adv 8: eabo4552

13. Gudipaty SA, Lindblom J, Loftus PD, Redd MJ, Edes K, Davey CF, Krishnegowda V, Rosenblatt J. (2017). Mechanical stretch triggers rapid epithelial cell division through Piezo1. Nature 543(7643):118–121. doi: 10.1038/nature21407.

14. Hannezo E, Heisenberg CP. (2019). Mechanochemical feedback loops in development and disease. Cell 178(1):12–25.

15. He L, Si G, Huang J, Samuel ADT, Perrimon N. (2018). Mechanical regulation of stem-cell differentiation by the stretch-activated Piezo channel. Nature 555(7694):103–106. doi: 10.1038/nature25744.

16. Jelínek R, Pexieder T (1970) Pressure of the CSF and the morphogenesis of the CNS. Foria Morphologica 18: 102–110.

17. Johansson PA, Dziegielewska KM, Ek CJ, Habgood MD, Liddelow SA, Potter AM, Stolp HB, Saunders NR (2006) Blood-CSF barrier function in the rat embryo. Eur J Neurosci 24:65–76

18. Johansson P, Dziegielewska K, Saunders N (2008) Low levels of Na, K-ATPase and carbonic anhydrase II during choroid plexus development suggest limited involvement in early CSF secretion. Neurosci Lett 442:77–80

19. Jones HC, Bucknall RM (1987) Changes in cerebrospinal fluid pressure and outflow from the lateral ventricles during development of congenital hydrocephalus in the H-Tx rat. Exp Neurol 98: 573–583

20. Kato S, Shindo A. Direct quantitative perturbations of physical parameters in vivo to elucidate vertebrate embryo morphogenesis. Curr Opin Cell Biol. 2024 Aug 24;90:102420.

21. Kier, WM (2012). The diversity of hydrostatic skeletons. J Exp Biol. 215, 1247–1257.

22. Landis, EM (1926) The capillary pressure in frog mesentery as determined by micro-injection methods. Am J Physiol 75: 548–570.

23. Li H, Tornberg J, Kaila K, Airaksinen MS, Rivera C. (2002) Patterns of cation-chloride cotransporter expression during embryonic rodent CNS development. Eur J Neurosci 16: 2358–2370.

24. Lun MP, Monuki ES, Lehtinen MK. Development and functions of the choroid plexus-cerebrospinal fluid system. Nat Rev Neurosci. 2015 Aug;16(8):445–57.

25. McCafferty RE, Wood ML, Knisely WH. Uterine contractions and intraamniotic pressures in gravid mice. Am J Obstet Gynecol. 90: 120–127, 1964.

26. Miyata T. Mechanical and physical interactions involving neocortical progenitor cells. In Neocortical Neurogenesis in Development and Evolution, Ed. Wieland Huttner. p. 119–136 Wiley 2023

27. Nelson CM, Gleghorn JP, Pang MF, Jaslove JM, Goodwin K, Varner VD, Miller E, Radisky DC, Stone HA. (2017) Microfluidic chest cavities reveal that transmural pressure controls the rate of lung development. Development 144: 4328–4335

28. Nourse JL, Leung VM, Abuwarda H, Evans EL, Izquierdo-Ortiz E, Ly AT, Truong N, Smith S, Bhavsar H, Bertaccini G, Monuki ES, Panicker MM, Pathak MM. Piezo1 regulates cholesterol biosynthesis to influence neural stem cell fate during brain development. J Gen Physiol. 154(10):e202213084. 2022

29. Okamoto M, Namba T, Shinoda T, Kondo T, Watanabe T, Inoue Y, Takeuchi K, Enomoto Y, Ota K, Oda K, Wada Y, Sagou K, Saito K, Sakakibara A, Kawaguchi A, Nakajima K, Adachi T, Fujimori T, Ueda M, Hayashi S, Kaibuchi K, Miyata T (2013). TAG-1-assisted progenitor elongation streamlines nuclear migration to optimize subapical crowding. Nat Neurosci 16:1556–1566

30. Olstad EW, Ringers C, Hansen JN, Wens A, Brandt C, Wachten D, Yaksi E, Jurisch-Yaksi N (2019) Ciliary Beating Compartmentalizes Cerebrospinal Fluid Flow in the Brain and Regulates Ventricular Development. Curr Biol 29: 229–241.e6.

31. Pathak MM, Nourse JL, Tran T, Hwe J, Arulmoli J, Le DT, Bernardis E, Flanagan LA, Tombola F. (2014). Stretch-actiated ion channel Piezo1 directs lineage choice in human neural stem cells. Proc Natl Acad Sci USA. 111(45):16148–53.

32. Pexieder T, Jelínek R (1970) Pressure of the CSF and the morphogenesis of CNS. II. Pressure necessary for normal development of brain vesicle. Physiol Bohemoslov 18: 181–192.

33. Praetorius J, Damkier HH (2017) Transport across the choroid plexus epithelium. Am J Physiol Cell Physiol 312:C673–C686

34. Rice R, Rice DPC, Olsen BR, et al. Progression of calvarial bone development requires Foxc1 regulation of *Msx2* and *Alx4*. Dev Biol. 2003;262(1):75–87.

35. Ruxton GD. The unequal variance t-test is an underused alternative to Student’s t-test and the Mann-Whitney U test. Behavior Ecol. 2006; 17: 688–690.

36. Shikata Y, Okada T, Hashimoto M, Ellis T, Matsumaru D, Shiroishi T, Ogawa M, Wainwright B, Motoyama J (2011) Ptch1-mediated dosage-dependent action of Shh signaling regulates neural progenitor development at late gestational stages. Dev Biol 349: 147–159

37. Tsujikawa K, Saito K, Nagasaka A, Miyata T (2022) Developmentall interdependent stretcher-compressor relationship between the embryonic brain and the surrounding scalp in the preosteogenic head. Dev Dyn 251: 1107–1122

